# Incorporating and compensating cerebrospinal fluid in surface-based forward models of magneto– and electroencephalography

**DOI:** 10.1101/037788

**Authors:** Matti Stenroos, Aapo Nummenmaa

## Abstract

MEG/EEG source imaging is usually done using a three-shell (3-S) or a simpler head model. Such models omit cerebrospinal fluid (CSF) that strongly affects the volume currents. We present a four-compartment (4-C) boundary-element (BEM) model that incorporates the CSF and is computationally efficient and straightforward to build using freely available software. We propose a way for compensating the omission of CSF by decreasing the skull conductivity of the 3-S model, and study the robustness of the 4-C and 3-S models to errors in skull conductivity.

We generated dense boundary meshes using MRI datasets and automated SimNIBS pipeline. Then, we built a dense 4-C reference model using Galerkin BEM, and 4-C and 3-S test models using coarser meshes and both Galerkin and collocation BEMs. We compared field topographies of cortical sources, applying various skull conductivities and fitting conductivities that minimized the relative error in 4-C and 3-S models. When the CSF was left out from the EEG model, our compensated, unbiased approach improved the accuracy of the 3-S model considerably compared to the conventional approach, where CSF is neglected without any compensation (mean relative error < 20% vs. > 40%). The error due to the omission of CSF was of the same order in MEG and compensated EEG. EEG has, however, large overall error due to uncertain skull conductivity.

Our results show that a realistic 4-C MEG/EEG model can be implemented using standard tools and basic BEM, without excessive workload or computational burden. If the CSF is omitted, compensated skull conductivity should be used in EEG.

## Introduction

The electrochemical activity of neurons gives rise to an electromagnetic field. In electro– and magnetoencephalography (EEG, MEG), the brain is studied by measuring this field on the scalp (EEG) or outside the head (MEG). These non-invasive remote measurements reflect the synchronous components of brain activity in a spatially blurred way. To interpret the measured fields in terms of neural sources and to understand the genesis of the signals, we need a forward model that gives a mathematical relationship between elemental sources and the resulting signals. A forward model consists of models for sources and volume conductor co-registered with a sensor model of the measurement; considering interpretation of measured signals in relation to neuronal sources, all these parts contain approximations or errors. This study focuses on volume conductor modeling in the context of experimental or clinical brain research.

The volume conductor model is based on physics of quasi-static electromagnetic field in conducting medium. The conductivity profile of the head is modeled based on description of head anatomy, and the resulting problem is solved for given source model using either analytical or numerical methods. The accuracy of the model depends on the level of anatomical detail, correctness of geometrical aspects and conductivity values, and implementation of the calculations. Computational techniques and roles of different tissues have in the case of EEG been evaluated in tens of studies; for a recent simulation study with guidelines, see [1]. Due to properties of magnetic fields in near-spherical geometry, MEG is considered less sensitive to modeling errors than EEG [2,3]. Consequently, MEG forward problem has not been as actively studied, although some detailed simulations exist [1,3,4].

Forward modeling in experimental source imaging is typically done using simple models that can be generated automatically or with minimal manual effort: MEG source modeling is carried out using either a spherically symmetric model or a single-compartment model shaped according to the inner skull boundary, and EEG modeling is done with a three-shell (3-S) model (brain, skull, scalp) that is either spherically symmetric or realistically shaped. Approximate conductivity boundaries for such a simple model are straightforward to segment and mesh from magnetic resonance (MR) images, and the resulting field computation problem is ideal for surface integral methods; in experimental work, the typical choice has been collocation-weighted boundary element method (BEM) [5,6]. Boundary-based modeling also enables easy visual evaluation of segmentation results and is computationally highly efficient. Starting from meshed boundary surfaces, an adequate 3-S BEM model and forward solution for the whole brain takes less than two minutes to build on a desktop computer [3].

In simulations or single-subject case studies, the forward model may contain over 10 tissue types [7]; also anisotropic tissues can be modeled [8]. Generating a model with such high detail demands extensive MRI acquisitions and/or computed tomography (CT) data, skillful use of many image processing tools, a lot of manual work, and advanced numerical solvers that typically apply finite element (FEM) or finite difference (FDM) methods. The gap between the most advanced simulation work and practical experimental use is thus large. Common arguments against the experimental use of high-detail models include 1) difficulties in obtaining a good MRI contrast between the most important tissue compartments (skin, skull, CSF, gray and white matter) and the resulting problems and geometrical errors in segmentation; 2) errors due to sensor co-registration; 3) errors due to unknown tissue conductivity values, especially the skull conductivity; 4) high computational cost, and 5) large amount of manual work by expert users using special tools. Issue 1 can be mitigated by acquiring both T1-weighted and T2-weighted MRI data to improve contrast between the cerebrospinal fluid (CSF) and skull, allowing automatic separation of CSF and inner skull boundary while at the same time having high contrast-to-noise ratio for segmenting the boundaries of gray and white matter. In this work, we will provide a solution to issues 4–5 and deepen understanding on issue 3.

Simulation studies have shown that highly conducting cerebrospinal fluid between the brain and skull has a major effect on EEG and a smaller effect on MEG [1,7,9]. The role of CSF has recently been illustrated also experimentally by measuring EEG in different head positions [10] and in a parametric-empirical-bayesian study that used a template head model and grand-average data from many subjects [11]. Segmentation pipeline and tools for realistic surface-based [12] and volume-based [13] four-compartment (4-C) models (brain, CSF, skull, scalp) using a combination of T1– and T2-weighted MR images have been recently presented, and toolkits for building and solving volume-based 4-C EEG and MEG models with FEM are available commercially (BESA, www.besa.de) and within academic community (NeuroFEM, [14]). For EEG, there is also a recently published FEM pipeline [15] and a surface-based toolbox that models simplified CSF, approximates skull from T1 images and provides layered 4-C BEM models [16]. In brief, among forward modeling community there is a consensus that the CSF should be included in at least the EEG models, and work is done to make that feasible for experimental researchers. Previous studies have, however, used model parameters and a comparison paradigm that are biased against the simplified models that omit the CSF. We propose that the omission of CSF should be compensated in model conductivities and that the effect of CSF should be assessed in these less biased circumstances.

## Compensating the omission of cerebrospinal fluid

Because the CSF has much better electric conductivity than the skull, the volume current coming from the brain tends to flow in the CSF instead of passing through the skull. One might intuitively think that a well-conducting compartment between the brain and scalp would amplify EEG, but as the CSF compartment provides a low-resistance (or, low field-energy) channel for the volume currents to return, the result is the opposite: the CSF between the brain and skull leads to overall weaker electric field and current density in the scalp than in the case of a direct contact between the brain and the skull. The layer-like part of CSF, shaped similar to the inner skull surface, thus attenuates scalp potentials and EEG signals; the effect of this layered part of CSF on EEG amplitudes is therefore similar to that of the poorly conducting skull layer. If the CSF is omitted from the model, lowering the skull conductivity can thus partially compensate the amplitude effects. As the smearing of scalp potentials depends on relative conductivities of soft tissues and skull and the conductivity contrast between the CSF and skull in a 4-C model is larger than that of the brain and skull in a 3-S model, this compensation should also decrease the overall errors in topography shapes. Because of the near-spherical shape of the head and different role of volume current in signal generation, the effect of CSF on magnetic field amplitude is more difficult to characterize: the layered part of the CSF that lies below the skull is supposed to have quite a small effect, depending on the variation of the thickness of CSF (if the head and CSF were spherically symmetric, it would have no effect at all), while the CSF in sulci guides volume currents to flow more radially towards (or away from) the skull, bend steeply to the layered part, and return through the same or another sulcus, producing potentially complex changes to the magnetic field.

In earlier studies, where the effect of CSF has been assessed, the comparison has been done between 4-C (or more detailed) and 3-S models so that the volume of CSF has in the 3-S model been labeled as brain, but the conductivities of other compartments have been kept the same as in the reference model. If we want to estimate, how well a 3-S model would do in experimental setting, this comparison is seriously biased against the 3-S model and does not yield the information of interest:

1. The omission of CSF has not been compensated for in the conductivities of the 3-S model.
2. The conductivity of the skull is implicitly assumed known, but in reality it has considerable uncertainty. The skull conductivity error affects both models, but if it is the only modeling error (like in the 4-C scenario of this work), it has larger relative effects than if it is only one of model errors, like in the 3-S approximation.
3. All boundary surfaces are assumed correct. The effect of a conductivity boundary on potential depends on the jump of conductivity across the boundary (see the integral equation for potential in subsection Physics and field computation). The boundary between the CSF and inner skull has the largest conductivity jump in the head, and in a model with CSF this jump is over five times as large as in a 3-S model. The pial boundary between the gray matter and CSF is very close to the sources, and the CSF compartment is thin deep in sulci and above some gyral crowns; a small shift of pial surface may change local volume conduction considerably.

In this paper, we address the issues #1 and #2. To study issue #3 in a realistic way, we would need segmentations and meshings generated with different MR (or CT) images and segmentation toolboxes like was done for 3-S and further simplified MEG models in [3]. As this work is, to our knowledge, the first detailed evaluation of surface-based 4-C models for MEG/EEG, we leave issue #3 for future work.

## Aims of the study

In this work, we 1) employ the SimNIBS pipeline [12] for making experimentally usable 4-C surface models for BEM; 2) verify the 4-C BEM, present a meshing guideline and share a fast, easy-to-use BEM toolbox; 3) propose and evaluate a method for compensating the omission of CSF by adjusting the skull conductivity in a 3-S model, and 4) evaluate the effects of numerical error, unknown skull conductivity, and omission of CSF, leading to expected error in geometrically correct 4-C and 3-S models.

## Methods and models

### Anatomical models

We built head models for two subjects (Head 1 and Head 2) using existing, previously shared anatomical data that had been collected at different imaging centers with different MRI scanners and protocols. The data were processed independently by different users using SimNIBS “mri2mesh” pipeline [12]. Two heads were needed in order to independently fit and test optimized and compensated conductivities. In addition, Head 2 was built using MR sequences different from those used by the SimNIBS team; thus the modeling of Head 2 serves also as an independent validation of the pipeline.

Head 1 was built based on boundary meshes of the publicly available SimNIBS example data, downloaded from http://www.simnibs.de. Written informed consent had been obtained, the protocol had been approved by the local ethics committee of Faculty of Medicine of University of Tübingen, and the data had been anonymized. Details on MR scans and segmentation process are available from the website or [12]. From this data set, we used the outer boundaries of white matter (*white*), gray matter (*pial*) and cerebellum, the inner and outer boundaries of the skull (*inner skull, outer skull*), and the outer scalp boundary (*scalp)*. The original surface meshes were slightly smoothed using “smoothsurf” function of iso2mesh toolbox [17] and a segmentation error at the scalp base was manually corrected.

Head 2 was built from a structural MRI dataset that had been collected for the NIH Human Connectome Project at Athinoula A. Martinos Center for Biomedical Imaging, Massachusetts General Hospital. The participants gave written consent, the data were anonymized, and the study protocol was approved by the institutional review board of the Partners HealthCare (Massachusetts General Hospital). The MRI data were acquired on the 3T Siemens MGH-USC Connectome scanner using a 64-channel RF receive coil array [18,19]. The basic structural dataset consists of a T1-weighted acquisition obtained at 1 mm resolution with multi-echo MPRAGE sequence and a T2-weighted acquisition obtained at 0.7 mm resolution with SPACE sequence. Basic MRI acquisition parameters are available in Connectome documentation

(http://www.humanconnectome.org/documentation/MGH-diffusion), and corresponding datasets and detailed acquisition parameters are publicly available through the Laboratory of Neuro Imaging Image Data Archive (LONI IDA, https://ida.loni.usc.edu) and the WU-Minn Connectome Database (ConnecomeDB, https://db.humanconnectome.org).

The segmentation and meshing for Head 2 was done using the same pipeline as for Head 1, with small modifications: First, the FreeSurfer [20–22] reconstruction was done using the T1-weighted data only as this was observed to result in slightly smoother boundary surface between the gray and white matter. Then, the SimNIBS processing pipeline “mri2mesh” was run employing both the T1-and T2-weighted acquisitions, with minor changes in parameters of the FSL brain extraction and surface generation tool “bet” [23,24]. Specifically, the “bet” command was run with clean-up option for the optic nerve and eye, and the basic brain extraction parameters were, based on visual inspection of MR scans and segmentation results, manually tuned to obtain accurate surfaces for skin, skull, and scalp. Finally, based on visual inspection of the results, surfaces were slightly smoothed employing FreeSurfer, meshfix [25], and iso2mesh tools. The resulting surfaces of inner skull and cerebellum intersected at skull base. We fixed this with a custom point-wise shifting routine that was applied also to the upper part of the cerebellum, ensuring a smooth gap between the cerebellum and pial surfaces.

For both heads, the basic anatomical modeling resulted in very dense triangle meshes for each segmented boundary. These meshes were re-triangulated using iso2mesh tool “meshresample” to various densities, specified according to the desired mean triangle-side length (TSL). The boundary of white and gray matter was tessellated at the TSL of 3 mm. This mesh was then used for the source space; the rest of the boundaries were used in the volume conductor model as conductivity boundaries. The conductivity boundaries close to the source space were meshed more densely than the more remote boundaries. In both heads, the gray matter was already in the dense meshes very thin in some regions (about 1% of the cortex was thinner than 1.5 mm; minimum thicknesses were below 0.5 mm). The thin regions were mainly in deep medial regions and in medial occipital cortex, but also in the posterior wall of central sulcus. These regions were automatically corrected by shifting the outer boundary of the white matter locally away from the pial surface, setting the minimum thickness of cortex to 1.5 mm in each model. The correction results in slightly different source spaces in different models, but the error due to the sub-millimeter shift of source position is negligible both in absolute scale and especially in comparison to the gained benefit in numerical accuracy in coarser models [3].

For both anatomical models, we built a 4-C reference model with dense meshes (TSL of 2 mm at the pial boundary, a total of 63354 vertices in Head 2) and five coarser 4-C models and corresponding 3S models. The meshings of each surface of Head 2 are described in Table 1, and example meshes are visualized in Fig 1. As we have a separate boundary for the cerebellum, our 4-C model actually consists of five sub-volumes and is not nested; we labeled both the cerebellum and the space inside the pial surface as brain.

**Fig 1.**
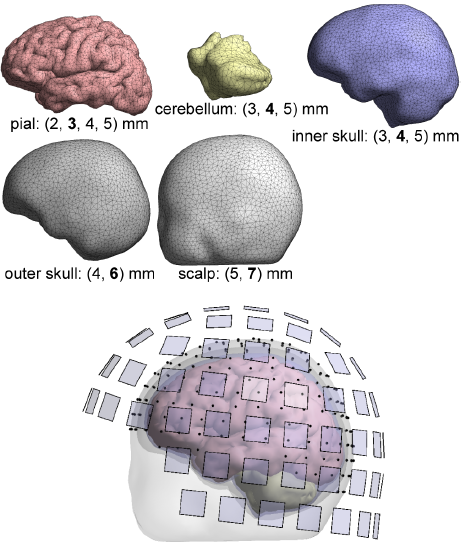
***Boundary meshes and the resulting four-compartment model***. *Below each boundary mesh, the used mesh densities in terms of mean triangle-side length TSL are listed; the TSL of the shown mesh is printed in boldface. EEG electrodes and magnetometer pickup coils are marked with black points and transparent blue squares, respectively*.

**Table I:**
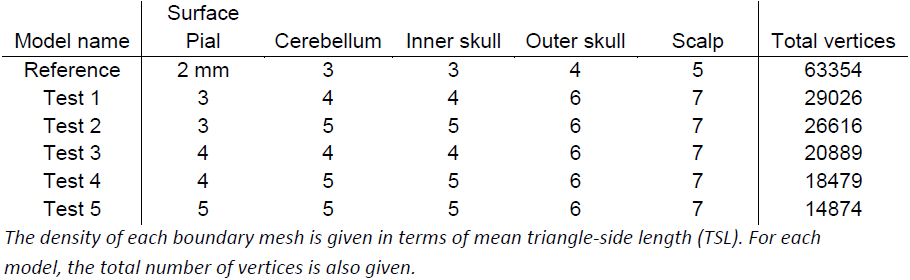
Mesh densities in reference and test models.

### Physics and field computation

EEG measures (the differences of) electric potential *Φ*, and MEG measures the flux of the magnetic field *B* due to all currents. The fields are modeled using standard quasi-static approximation of Maxwell’s equations [26]. The source activity is modeled in terms of primary current distribution 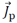 (e.g.,[6]) that gives rise to charge density that, in turn, creates a conservative electric field. The electric field drives volume currents in ohmic resistive medium. These definitions along with the law of charge conservation yield a Poisson-type equation 
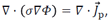

from which we can solve the electric potential *Φ* given primary current density 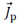 and conductivity σ. Magnetic field is generated by both primary and volume currents; in quasi-statics, we have the Biot-Savart law in the form 
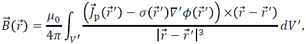

where – *σ∇Φ* is the volume current density, 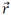 and 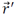 are the position vectors in field and integration spaces, and the integration space *V’* contains the whole volume conductor.

The conductivity profile of the head was modeled piecewise homogeneous and isotropic. Heading towards boundary-based methods, the Poisson equation is converted to surface-integral form using Green’s theorem and boundary conditions [6,27], leading to the surface-integral equation for potential 
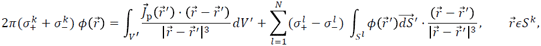

where 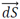 is the outward-directed different surface element, superscripts label boundary surfaces, subscripts + and – refer to compartments just outside and inside of the surface, and *N* is the number of boundary surfaces.

In a similar way, the contribution of volume currents to magnetic field in the Biot-Savart law is expressed as function of electric potential at conductivity boundaries, leading to Geselowitz equation for magnetic field [6,28],

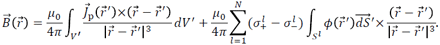

Then, the boundary potentials are discretized using some basis functions, and the method of weighted residuals (see [29]) is applied to the surface integral equation for potential. This discretization-and-weighting process is commonly referred to as the boundary element method (BEM).

We formulated the integral equations using the isolated source approach ISA [30,31]. We used linear basis functions and the accurate but computationally intensive Galerkin (LG) BEM [31,32] for the reference model, and both LG and less accurate but fast linear collocation (LC) BEM [5,6,29] for the test models. The magnetic field was computed using the Geselowitz equation, coupled to the BEM as described in [31]. Our field solvers are loosely based on the code of Helsinki BEM library [29], extended with the state-of-the-art ISA approach and Galerkin implementation described in [31] and further streamlined. The solvers have been previously verified [3,31,33].

For magnetic fields, we set up a sensor model that corresponds to 102 magnetometers of the 306-channel Elekta MEG system. The extent of pickup coils was taken into account by numerical integration over the coils (four quadrature points per magnetometer). For EEG, we used a 256-electrode ABC layout. Electrodes were not assumed to be in mesh nodes; the surface potential was interpolated from the mesh nodes to the electrode positions using the BEM basis functions. The sensor layouts are shown in Fig 1.

The conductivities for the soft tissues (brain, scalp) were chosen to commonly used 0.33 S/m. For CSF, we used 1.79 S/m [34]. The skull conductivity we express using soft tissue-skull resistivity ratio K; in the literature, estimates for K vary in the range of approx. 20-80, see [35]. For model verification and an example we chose K = 50, while for testing the effect of erroneous conductivity and assessing the error due to omitting the CSF we tested many values of K.

The electromagnetic field measured by EEG and MEG is understood to arise mainly from postsynaptic activity of pyramidal neurons in the cortex [2]. To model this activity, the primary current density was constrained to the cortex and discretized into a set of current dipoles that were placed into the vertices of the boundary of the gray and white matter and, following the main anatomical orientation of pyramidal dendrites, oriented perpendicular to that boundary. The orientation was taken from the original surfaces so it was the same in all models. For each constructed forward model, we computed MEG and EEG signal topographies for all elemental sources in the left hemisphere (in Head 2, 9350 separate dipoles in total). All analysis was based on these topographies.

### Metrics

We compared simulated topographies in sensor domain for each source separately using Relative Error (RE) and Correlation Coefficient (CC) metrics: 
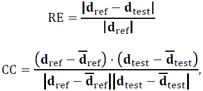
where **d**_ref_ and **d**_test_ are the reference and test topographies expressed in vector form and the overbar indicates the mean. The RE is a general difference measure that is sensitive to changes in both shape and amplitude of the topography, while CC characterizes the shape only: a small RE is achieved only if both shape and amplitude are similar, while two identically-shaped topographies have the CC of 1 regardless of the overall amplitude. Note that RE is normalized by the norm of the reference signal; the normalization can cause problems in the case of small **d**_ref_, as a difference that is small in absolute scale may be large in relative scale. This is seen in MEG comparisons as large RE values in weak topographies (quasi-radial and deepest sources) [3].

## Simulations and results

Before heading for quantitative assessment, we show an example that illustrates how the (uncompensated) CSF affects signal topographies and gives the reader an impression on the connection of RE and CC numbers to changes in signal patterns and amplitudes. Fig 2 shows two source positions and their corresponding MEG and EEG topographies in a 4-C and an uncompensated 3-S model. Both example sources produce strong signals in both modalities. The source marked in blue lies in the central sulcus, at the depth of 12 mm from the skull. The CSF has only a small effect on the shapes of MEG and EEG topographies, as shown by the high CC numbers. The amplitude of MEG topography changes less than the EEG amplitude. The source marked in red lies more superficially in the occipital cortex, at the depth of 6 mm. For this source, the CSF has larger effect on both the shape and amplitude of signal topography. For MEG, this source is an extreme case among strong sources; typically the effect of CSF is smaller. All subsequent analyses are carried out with similar computations of RE and CC between different models. In the following, we present results obtained in Head 2; conductivity optimizations were carried out independently in Head 1.

**Fig 2.**
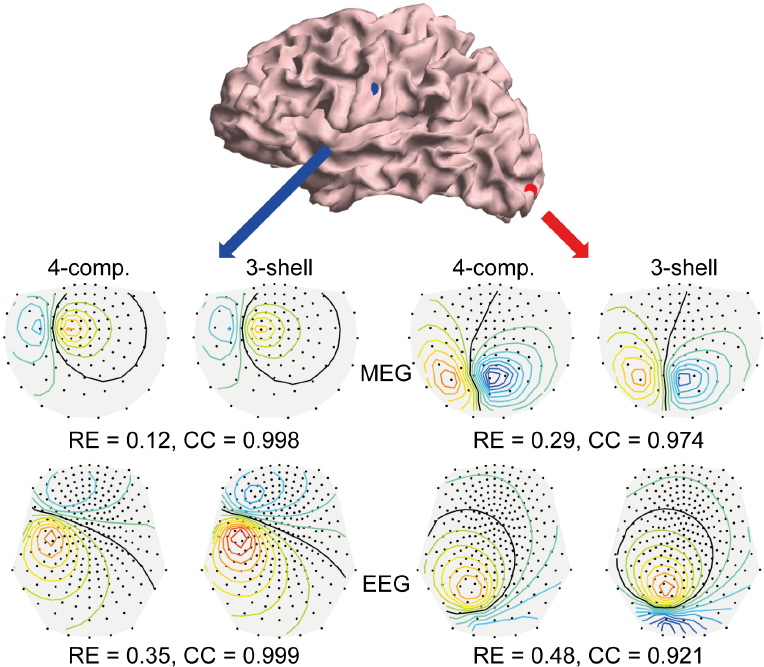
***Example on signal topographies and comparison metrics***. *At the top, two example source positions are shown on the cortex model. Below, the EEG and MEG topography maps computed for these sources using four-compartment (reference) and uncompensated three-shell (test) models are visualized; also the values for Relative Error RE and Correlation CC between the reference and test topographies are shown. All EEG plots are in the same scale as well as all the MEG plots. Contour lines have spacing of 0.1 times the maximum absolute value of the strongest topography; zero line is drawn in black, and yellow/red and blue mark positive and negative values, respectively*.

### Verification

To find out how sensitive the surface-based 4-C head model is to densities of boundary meshes and to the choice of BEM approach, we carried out a multi-resolution verification: We computed topographies for all sources using test models at various mesh densities (see Table 1) using both LG and LC BEM and compared them to our reference model (very dense mesh, LG BEM). The median and 16^th^ & 84^th^ percentiles (non-parametric equivalent to standard deviation) of the results in each test model are shown in Fig 3. In the case of MEG, the difference between the overall performance of LG and LC BEMs is very small, LG taking the lead as expected. The performances of test models 1 and 2 that have the TSL of 3 mm at the pial surface are very similar, and the same applied to models 3 and 4 where the pial surface is meshed at 4 mm TSL. For EEG, the results follow the same pattern and ranking as in MEG, but the errors are larger and there is a clear difference between the LG and LC BEMs. Overall, the error as function of both mesh density and chosen BEM type shows stable behavior: errors decrease when the mesh density increases or when the simple collocation BEM is replaced with the more accurate Galerkin BEM.

**Fig 3.**
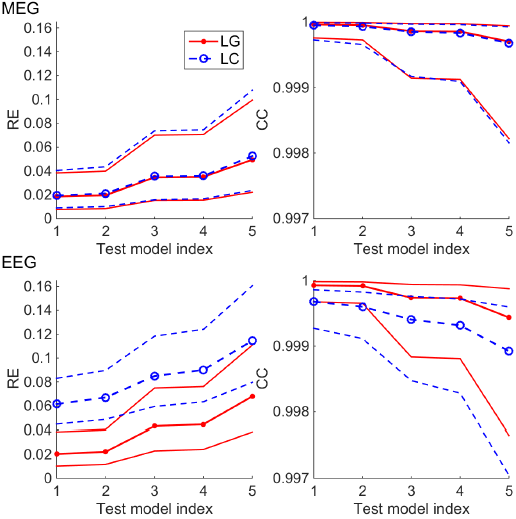
***Multi-resolution verification*** *of four-compartment head model in MEG (top row) and EEG (bottom row) using LG and LC BEM approaches (red and blue plots). The plots on the left side show the median and 16^th^ & 84^th^ percentiles of the Relative Error (RE) metric computed for all topographies, and the plots on the right show the corresponding plots of the Correlation (CC) metric*.

### Effect of skull conductivity and conductivity compensation

Next, we studied the role of skull conductivity in the 4-C model and tested, how well the omission of CSF can be compensated by adjusting the skull conductivity in the 3-S model. To focus on conductivity, all computations in this analysis were done in the dense reference geometry and the corresponding 3-S geometry using LG BEM. We built forward models with skull resistivity ratio K ranging from 20 to 80 (in steps of 10) in 4-C models and between 20 and 170 in 3-S models. The topographies of all sources were then compared across models that had different values of K. In the following, we label the K used in reference and test models with subscripts “ref” and “test”. For each pair or K_ref_ and K_test_, we took the median of the results across all sources. The results are shown in Fig 4.

**Fig 4.**
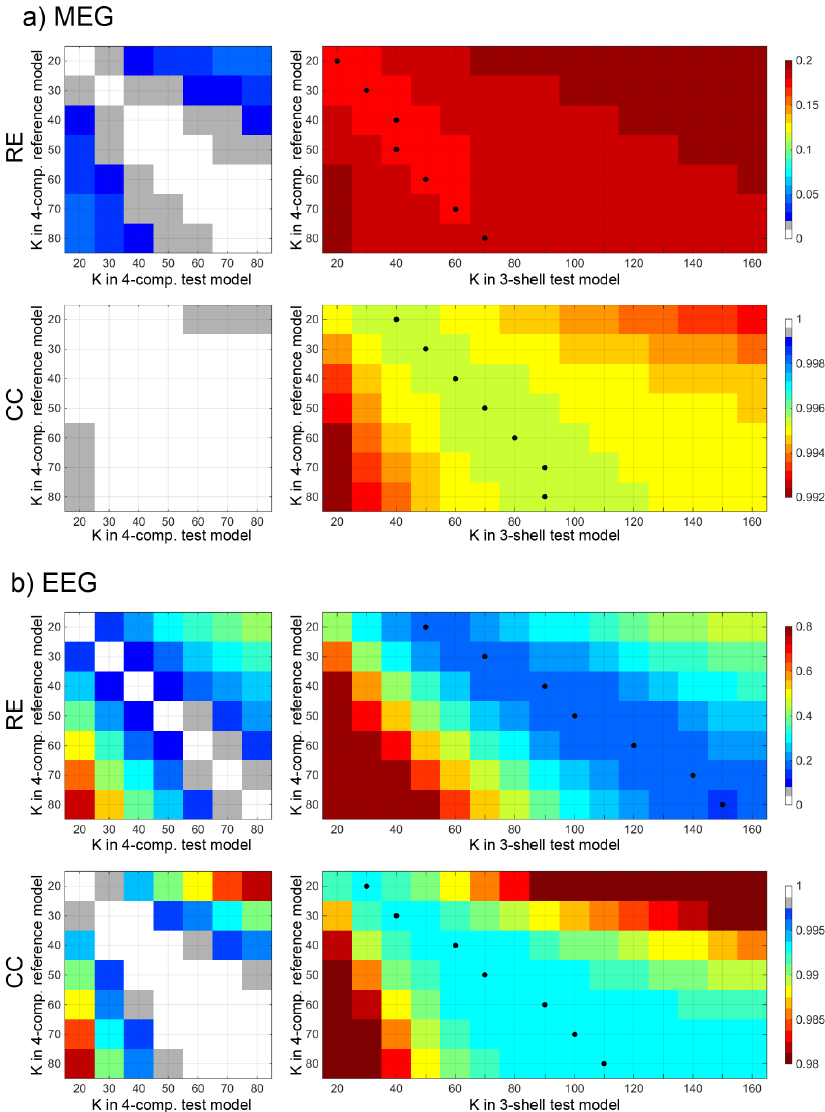
***Median RE and CC between the reference four-compartment model and test four-compartment (left) and three-shell (right) models as function of the skull resistivity ratio K*** *for a) MEG and b) EEG. The black dots show the best values for equivalent K in three-shell models. Note that color scales for MEG and EEG are not the same*.

For MEG, the RE due to erroneous skull conductivity ranges between 0% and 6% among 4-C models. In 3-S models, the combined effect of the skull conductivity and the omission of CSF is between 15% and 20%; the choice of K_test_ in a 3-S model thus has an effect of up to 5% (the values of K_test_ that produce smallest errors for each K_ref_ are plotted with black dots). In both 4-C and 3-S models, the K_test_ can be either over– or underestimated within a factor of about 1.5 without adding over 1%-unit of RE (as we specified K in steps of 10 and the effect depends also on the value of K_ref_, the quoted factors are rule-of-thumb approximations). The CC stays over 0.999 in all 4-C comparisons; only an extreme error in K_test_ drops the CC below 0.9995. In 3-S model, the CC is between 0.992 and 0.996; an extreme error in K_test_ drops CC below 0.994. Overall, the expected effect of mismatch of K_ref_ and K_test_ is small as long as the extreme ends of K range are avoided in the test model, and compensating the omission of CSF by changing K in the 3-S model has only a minimal effect.

In the case of EEG, the effect of skull resistivity ratio K on signal topographies is considerably higher than in MEG and also the compensation shows clear benefits: Among 4-C models, the choice K_test_ can lead to RE of up to 70%, if K_ref_ is large and K_test_ chosen small. If K_test_ is specified correctly within a factor of about 1.2 of K_ref_, the RE due to the mismatch of K stays below 10%, and having the K_test_ correct within a factor of 1.5 keeps the error below 20%. In 3-S models, the relative error RE is always over 15%. If compensation is omitted and the same K is used in the reference and 3-S test models, as has been done in earlier studies on the topic, the RE is between 45% and 50% for all values of K. The values of K_test_ that give best REs (between 15% and 20%) are about two to 2.5 times as large as the value of K_ref_; with small K_ref_, the omission of CSF is best compensated by using K_test_ of approx. 2.5 K_ref_, while with large K_ref_, the best compensation factor is roughly 2. Omission of compensation combined with underestimation of K_test_ may lead to very large RE in the 3-S model. The CC metric behaves overall in similar way to the RE metric, but the role of skull conductivity compared to the role of fourth compartment per se is smaller. Thus the 3-S model performs relatively worse in CC than in RE analysis. The 4-C model tolerates mis-specification of K_test_ by a factor of 1.5 with CC staying above 0.998, while the CC in the 3-S model is always below 0.994. The compensation shows clear benefits; the CC changes smoothly as factor of K_test_, the best compensation coefficient being about 1.5.

### Optimal and equivalent skull conductivity

Next, we analyze the data presented in the previous subsection further and answer an experimentally relevant question: If we know that the true skull conductivity ratio K_ref_ is within a certain range, how to choose the value of K_test_ in the 4-C and in the 3-S model? In addition, we assess, how large are the errors due to conductivity mismatch and due to omitting the CSF, when an optimized value of K is used.

As MEG is rather insensitive to the choice of K_test_, we defined the optimal values of K_test_ using EEG. To mimic a realistic situation for the 3-S model, we did not use any information from topographies of the corresponding 4-C model, but instead defined the optimal value in different head than in which we tested it. First, we computed the expectation value of RE and CC for each value of K_test_, when K_ref_ was varied between 20 and 80. Then, we chose the value of K_test_ that minimized the overall expected error. In the 4-C model this was straightforward, as both metrics had their best values at the same value of K. For the 3-S model, the RE metric behaved smoothly as function of K_test_, yielding three values of K that had REs within 1% of each other. Of these, we chose the one that was best in terms of CC. This analysis resulted in optimal K_test_ = K_opt_ = 50 for the 4-C model and optimal equivalent K_test_ = K_equi_ = 100 for the 3-S model. In the following, we use K_opt_ and K_equi_ in the test models.

Using K_opt_ in the 4-C model and K_equi_ in the 3-S model, we computed the overall errors due to erroneous skull conductivity and omission of CSF as the function of reference conductivity, still using the reference meshes and solver in all models; these results are shown in Fig 5. For MEG, the 4-C and 3-S models have the median RE of 0-3% and 17-18% and CC of over 0.999 and around 0.995, respectively. The variance of error as function of source position is larger in the 3-S than in the 4-C model. In the case of EEG, K_test_ has expectedly a large role: the median RE is between 0% and 29% in the 4-C model and between 18% and 33% in the 3-S model. The difference of errors between the 4-C and 3-S models decreases, as the value of K_ref_ deviates from 50 that was used as K_test_ in the 4-C model, illustrating issue #2 presented in Introduction. The median CC ranges between 0.991 and 1 for the 4-C model and 0.978 and 0.993 in the 3-S model. The CC of the 3-S model gets its maximum value at K_test_ = 70, off from the middle-of-the-range 50, because K_equi_ was defined based mainly on RE.

**Fig 5.**
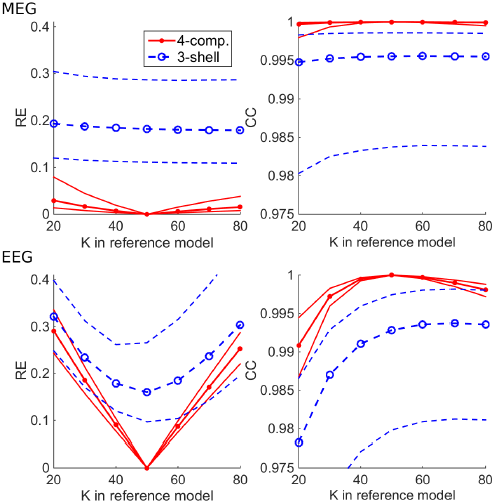
***Errors due to skull conductivity mismatch between the reference and test models***. *The median, 16^th^ percentile, and 84^th^ percentile of RE and CC metrics between the reference four-compartment (4-C) model and test 4-C and three-shell (3-S) models are plotted as function of the skull resistivity ratio K of the reference model. The 4-C and 3-S test models have the overall-best skull resistivity ratios of K_opt_ = 50 and K_equi_ = 100, respectively*.

### Expected error of four-compartment and three-shell models

In the final part of analysis, we answer a simple question: how large are the expected errors due to erroneous skull conductivity and the omission of CSF in 4-C and 3-S head models, when all geometrical information is correct and skull conductivity is known within a range?

We combined the effects of meshing and numerical error and the effects of unknown skull conductivity and omission of CSF. The mesh densities of the test models varied, and test models were built with optimized conductivity ratios (K_test_ = K_opt_ = 50 for the 4-C models, K_test_ = K_equi_ = 100 for the 3-S model) using both LG and LC BEMs. The differences between meshings of 3-S models were small compared to the overall error level of 3-S models, so we show results of one 3-S test model (that corresponds to model 4 of Table 1) only. The reference models were built using the reference meshes and LG BEM, varying K_ref_ between 20 and 80. For each K_ref_ and for all sources, we computed the RE and CC between the reference and test models and then took mean across all reference models.

Fig 6 shows, how the expected error behaves in 4-C models of different mesh densities and in a 3-S model with both BEM approaches. In all tests, models 1 & 2 and 3 & 4 performed equally well, as expected from verification computations (see Fig 3). For MEG, the median RE stayed below 6% for all 4-C models, while the 3-S model had the median RE of 19%. The overall accuracies of LG and LC BEM were almost identical. The median CC of all 4-C models was over 0.999, while the 3-S model was at 0.995. For EEG, the 4-C models had REs between 16% and 17%, and the 3-S model had 23%. The difference between LG and LC BEMs was small (much smaller than in the verification phase, see Fig 3). The median CCs of 4-S models were between 0.997 and 0.998, while the 3-S model gave the CC of 0.989.

**Fig 6.**
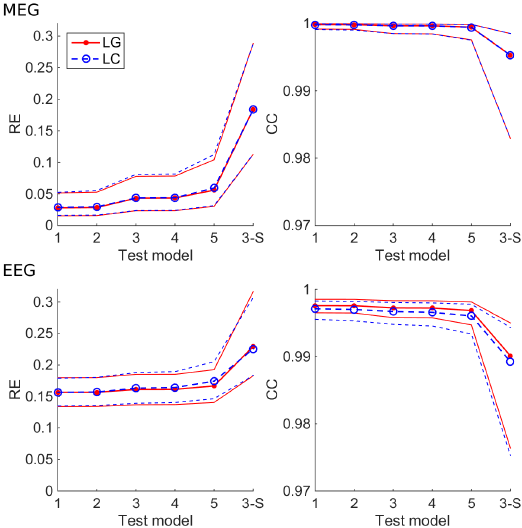
***Expected RE and CC of four-compartment (4-C) (models 1-5) and three-shell (3-S) test models, when the true skull resistivity ratio is known to be between 20 and 80***. *The* 4-C *test models have the skull resistivity ratio K = K_opt_ = 50, the 3-S test model has K = K_equi_ = 100. The plots show the median, 16^th^ percentile and 84^th^ percentile of the expected value of metrics for both LG and LC BEMs for different test models. Test models 1-5 are those described in Table 1, and model 3-S is the 3-S model that corresponds to test model 4*.

**Fig 6. Expected RE and CC of four-compartment (4-C) (models 1-5) and three-shell (3-S) test models, when the true skull resistivity ratio is known to be between 20 and 80**. *The* 4-C *test models have the skull resistivity ratio K = K_opt_ = 50, the 3-S test model has K = K_equi_ = 100. The plots show the median, 16^th^ percentile and 84^th^ percentile of the expected value of metrics for both LG and LC BEMs for different test models. Test models 1-5 are those described in Table 1, and model 3-S is the 3-S model that corresponds to test model 4*.

In Fig 7, we show the same expected errors as function of source position for representative 4-C and 3-S models (test model 4 of Table 1 and Fig 6 and the corresponding 3-S model, built using LC BEM). The results show that the largest errors in the 3-S EEG models occur at superficial sources close to tops of gyri, while the largest MEG errors occur both at gyrus tops and deep regions, especially at insula. Both EEG and MEG 3-S models have large errors at inferior frontal regions and in the anterior temporal lobe. Computation times for building these models in a standard PC that had 32 GB memory were about 11 min for the 4-C model and 90 s for the 3-S model. After building the models, EEG and MEG topographies for 10000 dipoles are then computed in 6 s in the 4-C model and 3 s in the 3-S model.

**Fig 7.**
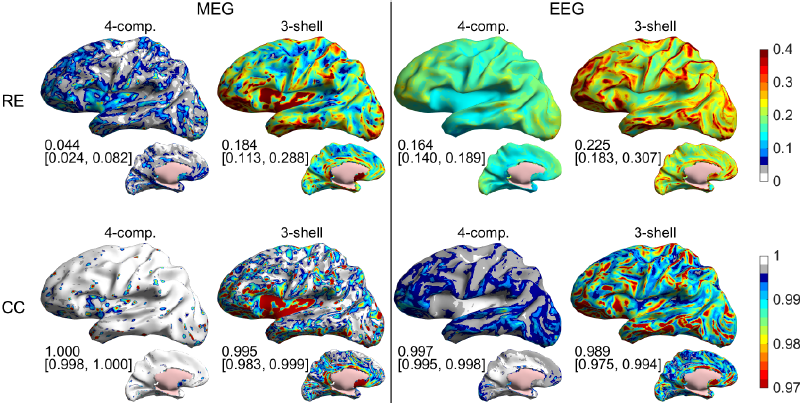
***The effect of numerical errors, conductivity errors, and the omission of CSF*** *: expected RE and CC of four-compartment (4-C) and three-shell (3-S) test models for all sources, when the true skull resistivity ratio is known to be between 20 and 80. Test models are built using the LC BEM. The* 4-C *test model (model 4 in Table 2 and Fig 6) has the skull resistivity ratio K = K_opt_ = 50, and the 3-S test model (the same as in Fig 6) has K = K_equi_ = 100. The numbers below the plots show median value and 16^th^ & 84^th^ percentiles of the plotted metrics. Results for MEG and EEG are shown on the left and side of the plot, respectively. The first row contains the RE results and the second row the CC results*.

***Fig 7. The effect of numerical errors, conductivity errors, and the omission of CSF**: expected RE and CC of four-compartment (4-C) and three-shell (3-S) test models for all sources, when the true skull resistivity ratio is known to be between 20 and 80. Test models are built using the LC BEM. The* 4-C *test model (model 4 in Table 2 and Fig 6) has the skull resistivity ratio K = K_opt_ = 50, and the 3-S test model (the same as in Fig 6) has K = K_equi_ = 100. The numbers below the plots show median value and 16^th^ & 84^th^ percentiles of the plotted metrics. Results for MEG and EEG are shown on the left and side of the plot, respectively. The first row contains the RE results and the second row the CC results*.

### Discussion

In this study, we constructed surface-based four-compartment (4-C) anatomical head models from MRI data using SimNIBS pipeline, built and verified MEG/EEG forward models with boundary element methods, presented and tested an approach for compensating the omission of CSF compartment by adjusting skull conductivity of a three-shell (3-S) model, and assessed the effects of mis-specified skull conductivity and omission of CSF on MEG/EEG signal topographies. The main results are:

1. A 4-C MEG/EEG model is straightforward to build using SimNIBS pipeline and standard BEM techniques, assuming good-quality T1-and T2-weighted MRI scans.
2. Simple and fast LC BEM (formulated with isolated-skull approach) is adequate for making 4-C models for experimental MEG/EEG use. With this BEM approach, we recommend meshing the pial surface, inner skull boundary, cerebellum, outer skull boundary, and scalp at triangle-side lengths of 4 mm, 5 mm, 5 mm, 6 mm, and 7 mm or less, respectively, and ensuring that dipolar sources are at least 1.5 mm from the pial surface. The building of such an LC ISA model takes about 11 minutes on a standard PC.
3. The omission of CSF can in 3-S EEG model be partially compensated by decreasing the skull conductivity by a factor of 1.5-2.5. The relative error of the compensated model is less than half of the error of a conventional, uncompensated model. The MEG model does not essentially benefit from compensation.
4. Even with a compensated model, the effect of omission of CSF on signal topographies is much larger than the errors due to coarse meshing and numerical approximations. In EEG, also the choice of skull conductivity has a larger influence than meshing and numerical accuracy do.
5. If the true skull resistivity ratio K is assumed to be between 20 and 80, the optimal choice for K in a 4-C model is 50 and in a 3-S model 100. In this case, the expected relative errors for signal topographies due to mis-specified skull conductivity and omission of CSF in 4-C and 3S MEG models are 4% and 18%, and in corresponding EEG models 16% and 23%, respectively, when the reference model has four compartments and is assumed geometrically correct. Corresponding correlation coefficients are 1 and 0.995 for MEG and 0.997 and 0.989 for EEG. Thus, the omission of CSF has larger overall effect on MEG than on EEG (as measured by RE), but the effect of omission of CSF on shapes of signal topographies is larger in EEG (as measured by CC).

#### Anatomical modeling

Our anatomical models were generated with SimNIBS pipeline that uses FreeSurfer, FSL, and meshfix toolboxes. All these tools are freely available to academic community. FreeSurfer is an extensively validated and documented standard tool in anatomical MR analysis in experimental brain research. In our experience, most laboratories that use individual realistically shaped head models in MEG or EEG analysis have FreeSurfer pipelines and compatible T1-weighted MRI sequences available. FSL is also widely used and freely available. Combining the results obtained with FreeSurfer and FSL and applying mesh manipulation tools such as those offered by “meshfix” to produce geometrically correct surfaces requires, however, some expertise and experience. For experimental use, the easy-to-use wrapper provided by SimNIBS thus greatly facilitates the combination and use of all these tools. For an experimentally oriented neuroimaging researcher, this modeling approach should be feasible to adopt, assuming that the pipeline has been set up and adjusted to the MRI scanner and the imaging protocols available.

While it is possible to employ only T1-weighted images in FreeSurfer/FSL segmentation of the brain and non-brain tissue compartments, using both T1 and T2 data is expected to yield better results at the cost of the added scan time. For present MRI systems this cost is not particularly drastic as fast high-resolution protocols are available for both T1 and T2 weighted acquisitions (the total acquisition time for Head 2 was approx. 12 minutes). To avoid problems in the segmentation of the skull, particular attention should be paid to the chemical shift between the fat and water in fatty tissue (e.g., the dura, spongy bone of skull, and skin). The bandwidth should be matched between the T1 and T2 acquisitions, and it should be high enough to avoid excessive blurring/shifting of the tissue boundaries due to displaced fat signal while maintaining sufficient SNR. If this is not feasible with a single acquisition, high-bandwidth scans can be used in conjunction with low-bandwidth scans to mitigate the SNR loss [12]; for Head 2 this was, however, not necessary.

The MRI data of Head 2 were obtained with a 3T Siemens “Connectom” scanner that is customized for high-performance diffusion imaging and equipped with an in-house built 64-channel RF head receive coil. In this study, only conventional structural data are needed and similar anatomical scans could be readily obtained using, for instance, a Siemens 3T Skyra scanner with commercially available 32-channel receive head coil. Finding the optimal combination of imaging parameters and FreeSurfer/FSL options requires some experimentation when setting up the imaging pipeline, but this work pays off in smaller manual workload in actual use. For best results, parameters of the FSL brain extraction tool “bet” need to be adjusted for the specific scanner and imaging parameters. This, again, may require some work in the setup phase.

As the SimNIBS pipeline uses different toolboxes for different parts of the model, some attention and post-processing may be needed. While the pipeline ensures that pial surface is fully inside the skull, in our case in Head 2 the cerebellum was intersecting at the skull base. We corrected this using the same custom tool that we used for adjusting part of the white/gray surface. Once the imaging and segmentation pipeline is running smoothly, the additional interactive workload of making a 4-C model instead of making a FreeSurfer-based single-or 3-S model is of the order of one hour per subject. As usual, the resulting segmentations and surfaces should be carefully inspected and, if necessary, manually corrected before proceeding with the MEG/EEG field computations.

#### Field computation

We carried out the field modeling using two types of boundary element methods, formulated using isolated source approach (variant #2 of [31]). From previous 3-S studies [32,33] it was known that the Galerkin BEM (LG) is in idealized 3-S EEG models more accurate than the collocation BEM (LC); our results expectedly confirmed that this is the case also in 4-C models. More interestingly, our results showed that the difference between the LG and LC methods in a corresponding MEG modeling scenario is small. When typical uncertainty is added to the model, the overall errors of LG and LC methods are comparable also in EEG, as the additional error due to the LC discretization is much smaller than the role of conductivity error or especially the role of omitting the CSF. For experimental use of 4-C or 3-S model in, e.g., source imaging, the LC BEM is thus sufficient. For computational analysis of high-detail models or for making reference models in method development, we recommend the LG approach.

The collocation BEM is commonly considered unreliable, when sources are close to the skull (see [33]). In the case of CSF, the situation is, however, different: the layer of CSF between the brain and the skull smooths the electric field and thus facilitates discretization of the potential at the inner skull boundary; the insensitivity of the 4-C model to the skull meshing is seen in results shown in Figs 3 and 6. The effects of poorly-known skull conductivity further compensate the numerical errors of collocation BEM, leading to similar expected accuracies for the LC and LG BEMs in an experimental situation, given appropriate meshing of the pial surface (in our case, 4 mm triangle-side length) and source depth. In this work, we ensured 1.5 mm distance between the sources and the pial boundary, slightly shifting sources that were too close to the boundary. As we did not model the white-matter anisotropy or divide the brain to compartments of gray and white matter, and also the brain conductivities are not well known, this kind of small shifting is safe as long as the orientations are kept unchanged. Other ways for ensuring numerical stability would be either to mesh the pial surface more densely in thin regions of the cortex, or to use continuous, distributed source representation instead of dipoles [36]. We could also approximate extended dipoles using a pair or set of spaced monopoles, as is done in volume-based approaches. In verification phase, we tested the continuous model, representing elemental sources with linear hat functions spanned on the white/gray boundary. Verification results (not shown) were systematically but only minimally better, so we did not study this computationally intensive approach further. It may, however, become valuable, if sources need to be placed closer to the pial surface due to, for example, including the white matter as conductivity boundary, or if fields are computed close to the sources, for example, in the case of modeling of electrocorticographic signals.

#### Compensating the omission of CSF

In earlier studies on the topic, the effect of inclusion or omission of CSF or any other compartment in a multi-compartment model has been assessed in a biased way: the CSF has just been left out and replaced with brain tissue, without adjusting other conductivities. In some works, this has correctly been called “neglecting” the CSF. Our compensation approach, in which the skull conductivity of the 3-S model is adjusted, is based on simple physics-based reasoning, and the actual compensation coefficient is estimated using a more detailed model. Thus, our compensated 3-S model is an informed attempt to get the best out of the simple 3-S model after making a decision to consciously leave the CSF out; this justifies our use of verb “omit”. We fitted the compensation coefficient using information on Head 1, and we used it in Head 2; thus our assessment results shown in Figs 5–7 are not biased in the favor of the simplified model.

To our knowledge, this is the first work, where conductivity outside the omitted compartment is adjusted in a simplified model. Dannhauer et al. [37] simulated the effect of skull fine-structure (highly resistive compact bone and less resistive spongy bone) on EEG; they estimated that for compact– and spongy-bone conductivities of 0.0064 S/m and 0.02865 S/m, the overall optimal conductivity for a homogeneous skull model was about 1.5 times the conductivity of the compact bone. Also the conductivity of scalp can be interpreted as a joint equivalent of conductivities of muscle and fat, as was done in [38] even though we are not aware of detailed simulations on that topic.

Another way for compensating the omission of CSF would be to increase the conductivity of the brain compartment in the 3-S model. Also this approach is physically motivated, by matching the average work done by the electric field in driving current in the intracranial spaces of 4-C and 3-S models. We ran a simple test on this, using Head 2, anatomical model 4, with skull resistivity ratio K_ref_ = K_test_ = 50, and found that increasing the conductivity of the brain compartment in the 3-S model by a factor of 1.5 reduced the median RE in EEG roughly as well as the adjustment of the skull conductivity, but had practically no effect on CC. This suggests that improved compensation performance should result from optimizing the CC by adjusting the skull conductivity and then tuning the amplitude by slightly increasing the brain conductivity. Indeed, decreasing the skull conductivity by a factor of 1.5 (see Fig 4b) and increasing the brain conductivity by a factor of 1.25 resulted in the case of EEG in a slightly better combination of RE and CC than the compensation done by adjusting the skull conductivity only. To properly assess the effect of brain conductivity on topographies, we would also need to assess the variation and errors in brain conductivity; this would mean adding an extra dimension to optimization and all conductivity-related analyses. We leave detailed analysis of multi-parameter compensation and role of brain conductivity for future studies.

#### Role of CSF in MEG and in EEG

The results shown in Figs 4–7 indicate that the omission of CSF causes larger overall error on MEG than on EEG, both in absolute and relative terms. The effect of omission of CSF on shapes of signal topographies is, however, smaller for MEG than for EEG. The latter result means that if there are one or a couple of focal sources that produce strong signals (say, a somatotopic experiment), the omission of CSF would overall disturb the source localization less in MEG than in EEG. On the other hand, incorporating the CSF might benefit MEG more than EEG in a more general and complex analysis task, where the use of strong priors such as focality of the sources is not warranted.

Even though the added overall RE due to omitting the CSF is smaller in EEG than in MEG, there is a feature in the spatial error distributions that stresses the importance of CSF in EEG: the 3-S model produces large errors for sources at tops of gyri (Fig 7). These are the regions of highest sensitivity for EEG and typically low sensitivity for MEG. Thus, the error in the 3-S EEG model may both mask weaker sources and increase both spread from and cross-talk to these regions, corrupting the EEG analysis. In MEG, this problem would only arise, if the source estimation algorithm amplifies weak sources in an uncontrolled or non-robust way.

Our results were obtained using all cortical sources. Compared to EEG, MEG has strong orientation selectivity (pseudo-radial sources produce weak MEG signals) and the amplitude of signals decays faster as function of source depth. Our source space contains thus areas, whose MEG signals would in real-world conditions be buried under noise. We did not leave these weakest sources out, like was done in the main analysis of [3]: As we deal with both MEG and EEG here, leaving the weak-MEG sources out would bias the comparisons. More importantly, the choice of sources to be omitted is based on the forward model and is thus sensitive to model simplifications and errors, and in many applications, signal averaging can considerably enhance signals from low-gain sources. On the other hand, we often have known region-of-interests or prior information on source location, and if this information points to deep cortical regions, MEG might not be the tool chosen for the task. We thus tested, how our MEG results would change, if weak sources are omitted like was done in [3]: the RE and CC values improve slightly, especially in the 3-S model; the shapes, overall levels and rankings of result plots stay the same, and the benefit of modeling the CSF remains clear. In analysis of Fig 7, the median RE in the 4-C model drops from 0.044 to 0.039, and in the 3-S model from 0.184 to 0.169; the median CC in the 3-S model increases from 0.995 to 0.997. Comparing to corresponding numbers for EEG, a 3-S MEG model is with strong sources roughly as accurate as a 4-C EEG model. As MEG topographies are in general less correlated than EEG topographies, this means that MEG and a 3-S model can likely resolve activities of different strong sources better than EEG and a 4-C EEG model can.

#### Future directions

In this work, we studied thoroughly the connection between CSF compartment and modeled skull conductivity. Plots in Fig 4 show that compensating the omission of CSF by adjusting the skull conductivity approximately halves the overall error when compared to a conventional modeling situation, where the CSF is just left out. In our analysis on the role of unknown skull conductivity, the skull conductivity was optimized in both 4-C and 3-S test models using as much same information as possible, avoiding bias. Future studies that aim at recommendations regarding models used in experimental research should follow similar logic: When high-detail models are presented and evaluated, high-detail simulations should also be used for tuning the parameters of the simplified models towards optimal. If the role of an additional model compartment is studied globally in an anatomically normal head assuming the conductivities and geometries error-free, the simplified model can be optimized in the same head geometry where it is assessed. If many head geometries are available, the best way would be to use a learning set for defining optimal and equivalent conductivities and testing these against many values of reference conductivity in an independent test set. Before heading for laborious group simulations using high-detail models, one should, however, consider errors in conductivities and in model geometry.

In this study, we varied the skull conductivity and assumed other conductivities known. In the future, also other conductivities should be evaluated. For example, the conductivity used in homogeneous brain model should be optimized in a model that contains both gray and (preferably anisotropic) white matter; only then can the effect of omitting the distinction between the gray and white matter be fairly evaluated. Further, strong conclusions should not be drawn without assessing also the effect of conductivity errors—estimated values for the conductivity of gray matter vary between 0.1 S/m [39] and 0.33 S/m [40]. Similar comparisons should be done for the scalp, in which the recently-used conductivities range between 0.25 S/m [38] and 0.43 S/m [1]. Electric impedance tomography carried out at low frequencies using EEG electrode setup is a promising way for specifying the equivalent scalp and skull conductivities of a 3-S or 4-C model; see [41].

In addition to conductivity, also differences and errors in model geometries should be evaluated. Such a study could be done by segmenting the same head using different software toolboxes and MR sequences and building forward models using different (verified) techniques, like was done for 3-S MEG models in [3]. In such a study, avoiding the bias in the choice of conductivities is essential. Another, more controlled approach would be to artificially perturb the boundary surfaces; this kind of study was recently done in the case of transcranial magnetic stimulation [42]. Studies should extend also to anatomically abnormal heads, for example those containing CSF-filled lesions. Regarding experimental use, detailed assessment of the effects of geometrical errors and approximations in the skull model is also an interesting future topic: While current high-detail approaches use T2-weighted MR sequences, standard anatomical brain imaging is carried out using T1-weighted sequences that have poor image contrast between the compact skull bone and CSF. Thus, statistical or rule-based algorithms, like that used in [16], for estimating the skull from T1-weighted images, should be developed and independently evaluated.

We did not do source estimation computations in this work, as we wanted to do general analysis on forward models, and results obtained with a specific estimation technique generalize poorly to other techniques. For example, the value of skull conductivity of a 3-S model is considered important in EEG single-dipole localization, but in linear distributed source estimation it plays only a small role [35]. Source estimation algorithms use the measured data, forward model, and prior information in varying ways, so the best way to understand the effect of forward-model errors and simplifications on the performance of a specific estimation technique is to set up a simulation for testing exactly that technique. Correspondingly, when new inverse modeling approaches are proposed or existing techniques compared, they should be tested for robustness to common model errors such as the omission of CSF.

Our analysis was done using MEG and EEG. Due to the reciprocity principle [43], the modeling approach presented here can also be applied to forward problems of transcranial magnetic stimulation (TMS) [44] and transcranial direct current and alternating-current stimulation (TDCS, TACS) [12,38]. In those problems, coils and electrodes are larger, and fields are thus expected to behave more smoothly than in the MEG/EEG problem studied in this work. On the other hand, compared to MEG sensor array, the TMS coil is more local and closer to the scalp; the role of volume conductor thus depends on coil position. Specific studies on MEG, TDCS, and TACS are thus needed to optimize the level of detail of head and sensor models.

Last, the effect of model detail and chosen conductivities should be tested in experimental studies. To make generalizable conclusions or recommendations from experimental data, we need studies that focus on different brain regions and that apply different kinds of prior information and source estimation techniques, from simple dipole localization and beamforming to sparse distributed models and multiple-region connectivity analysis. To facilitate further studies and experimental use, our LCISA BEM solver is available from the corresponding author for academic use.

### Conclusions

In this work, we presented and verified a 4-C BEM model for MEG and EEG, suggested a technique for compensating the omission of CSF in a 3-S model, and assessed the role of skull conductivity and the effect of omitting the CSF. Using SimNIBS pipeline for anatomical modeling, a 4-C model is straightforward to build. For experimental use, basic collocation BEM is adequate. We recommend 4-mm triangle size for the pial mesh; inner and outer skull and scalp meshes can be tessellated with 5-mm, 6-mm, and 7-mm triangles, respectively. Computation time for building such a BEM model on a PC is approximately 11 min.

If the CSF is not modeled, the skull conductivity of the 3-S EEG model should be decreased by a factor of approx. 2 in order to compensate the omission of CSF. If skull-soft tissue resistivity ratio K is assumed to lie between 20 and 80, good choices for K in 4-C and 3-S models are K = 50 and K = 100, respectively. As MEG is rather insensitive to the choice of conductivity values, it also does not benefit from compensation, and skull conductivity does not need to be adjusted.

The omission of CSF has roughly as large effects on MEG and compensated EEG. Due to its sensitivity to skull conductivity, EEG has in the 4-S model considerably higher expected error than MEG has, when a set of dense 4-S models with varying skull conductivities are used as reference.

The collocation BEM solver used in this work is available for academic use from the corresponding author.

## Acknowledgments

The calculations presented in this work were performed using computer resources within the Aalto University School of Science “Science-IT” project. The study was partially funded by the National Institutes of Health under NIBIB award number R00EB015445. The funding source had no role in the conduct or reporting of the research. The original data for Head 2 were collected under National Institutes of Health Blueprint Initiative for Neuroscience Research grant U01MH093765.

## References

1. Vorwerk J, Cho JH, Rampp S, Hamer H, Knosche TR, Wolters CH. A guideline for head volume conductor modeling in EEG and MEG. Neuroimage. 2014;100: 590–607.

2. Hamalainen MS, Hari R, Ilmoniemi RJ, Knuutila J, Lounasmaa OV. Magnetoencephalography— theory, instrumentation, and applications to noninvasive studies of the working human brain. Rev Mod Phys. 1993;65: 413.

3. Stenroos M, Hunold A, Haueisen J. Comparison of three-shell and simplified volume conductor models in magnetoencephalography. Neuroimage. 2014;94: 337–348.

4. Ramon C, Haueisen J, Schimpf PH. Influence of head models on neuromagnetic fields and inverse source localizations. Biomed Eng Online. 2006;5: 55.

5. de Munck JC. A linear discretization of the volume conductor boundary integral equation using analytically integrated elements. IEEE Trans Biomed Eng. 1992;39: 986–990.

6. Sarvas J. Basic mathematical and electromagnetic concepts of the biomagnetic inverse problem. Phys Med Biol. 1987;32: 11–22.

7. Ramon C, Schimpf P, Haueisen J, Holmes M, Ishimaru A. Role of soft bone, CSF and gray matter in EEG simulations. Brain Topogr. 2004;16: 245–248.

8. Güllmar D, Haueisen J, Eiselt M, Giessler F, Flemming L, Anwander A, et al. Influence of anisotropic conductivity on EEG source reconstruction: investigations in a rabbit model. IEEE Trans Biomed Eng. 2006;53: 1841–1850.

9. Akalin Acar Z, Makeig S. Effects of forward model errors on EEG source localization. Brain Topogr. 2013;26: 378–396.

10. Rice JK, Rorden C, Little JS, Parra LC. Subject position affects EEG magnitudes. Neuroimage. 2013;64: 476–484.

11. Strobbe G, van Mierlo P, De Vos M, Mijovic B, Hallez H, Van Huffel S, et al. Bayesian model selection of template forward models for EEG source reconstruction. Neuroimage. 2014;93: 11–22.

12. Windhoff M, Opitz A, Thielscher A. Electric field calculations in brain stimulation based on finite elements: an optimized processing pipeline for the generation and usage of accurate individual head models. Hum Brain Mapp. 2013;34: 923–935.

13. Lanfer B. Automatic Generation of Volume Conductor Models of the Human Head for EEG Source Analysis. Doctoral Thesis, University of Munster. 2014.

14. SimBio Development Group. SimBio: A generic environment for bio-numerical simulations. 2015.

15. Ziegler E, Chellappa SL, Gaggioni G, Ly JQ, Vandewalle G, Andre E, et al. A finite-element reciprocity solution for EEG forward modeling with realistic individual head models. Neuroimage. 2014;103: 542–551.

16. Akalin Acar ZA, Makeig S. Neuroelectromagnetic forward head modeling toolbox. J Neurosci Methods. 2010;190: 258–270.

17. Fang Q, Boas D. Tetrahedral mesh generation from volumetric binary and gray-scale images. Proc IEEE Int Symp Biomed Imaging 2009. 2009. pp. 1142–1145.

18. Keil B, Blau JN, Biber S, Hoecht P, Tountcheva V, Setsompop K, et al. A 64-channel 3T array coil for accelerated brain MRI. Magn Reson Med. 2013;70: 248–258.

19. Setsompop K, Kimmlingen R, Eberlein E, Witzel T, Cohen-Adad J, McNab JA, et al. Pushing the limits of in vivo diffusion MRI for the Human Connectome Project. Neuroimage. 2013;80: 220–233.

20. Dale AM, Fischl B, Sereno MI. Cortical surface-based analysis. I. Segmentation and surface reconstruction. Neuroimage. 1999;9: 179–194.

21. Fischl B, Sereno MI, Dale AM. Cortical surface-based analysis. II: Inflation, flattening, and a surface-based coordinate system. Neuroimage. 1999;9: 195–207.

22. Fischl B. FreeSurfer. Neuroimage. 2012;62: 774–781.

23. Smith SM. Fast robust automated brain extraction. Hum Brain Mapp. 2002;17: 143–155.

24. Jenkinson M, Pechaud M, Smith S. BET2: MR-Based Estimation of Brain, Skull and Scalp Surfaces. Eleventh Annual Meeting of the Organization for Human Brain Mapping. 2005.

25. Attene M. A lightweight approach to repairing digitized polygon meshes. Vis Comput. 2010;26: 1393–1406.

26. Plonsey R, Heppner DB. Considerations of quasi-stationarity in electrophysiological systems. Bull Math Biophys. 1967;29: 657–664.

27. Barnard AC, Duck IM, Lynn MS. The application of electromagnetic theory to electrocardiology. I. Derivation of the integral equations. Biophys J. 1967;7: 443–462.

28. Geselowitz DB. On the magnetic field generated outside an inhomogeneous volume conductor by internal current sources. IEEE Trans Mag. 1970;6: 346—347.

29. Stenroos M, Mäntynen V, Nenonen J. A Matlab library for solving quasi-static volume conduction problems using the boundary element method. Comput Methods Programs Biomed. 2007;88: 256–263.

30. Hamalainen MS, Sarvas J. Realistic conductivity geometry model of the human head for interpretation of neuromagnetic data. IEEE Trans Biomed Eng. 1989;36: 165–171.

31. Stenroos M, Sarvas J. Bioelectromagnetic forward problem: isolated source approach revis(it)ed. Phys Med Biol. 2012;57: 3517–3535.

32. Mosher JC, Leahy RM, Lewis PS. EEG and MEG: forward solutions for inverse methods. IEEE Trans Biomed Eng. 1999;46: 245–259.

33. Stenroos M, Nenonen J. On the accuracy of collocation and Galerkin BEM in the EEG/MEG forward problem. Int J Bioelectromagn. 2012;14: 29–33.

34. Baumann SB, Wozny DR, Kelly SK, Meno FM. The electrical conductivity of human cerebrospinal fluid at body temperature. IEEE Trans Biomed Eng. 1997;44: 220–223.

35. Stenroos M, Hauk O. Minimum-norm cortical source estimation in layered head models is robust against skull conductivity error. Neuroimage. 2013;81: 265–272.

36. von Ellenrieder N, Valdes-Hernandez PA, Muravchik CH. On the EEG/MEG forward problem solution for distributed cortical sources. Med Biol Eng Comput. 2009;47: 1083–1091.

37. Dannhauer M, Lanfer B, Wolters CH, Knösche, T. R. Modeling of the human skull in EEG source analysis. Hum Brain Mapp. 2011;32: 1383–1399.

38. Opitz A, Paulus W, Will S, Antunes A, Thielscher A. Determinants of the electric field during transcranial direct current stimulation. Neuroimage. 2015;109: 140–150.

39. Akhtari M, Mandelkern M, Bui D, Salamon N, Vinters HV, Mathern GW. Variable anisotropic brain electrical conductivities in epileptogenic foci. Brain Topogr. 2010;23: 292–300.

40. Geddes LA, Baker LE. The specific resistance of biological material-a compendium of data for the biomedical engineer and physiologist. Med Biol Eng. 1967;5: 271–293.

41. Dabek J, Kalogianni K, Rotgans E, van der Helm F, Kwakkel G, van Wegen E, et al. Determination of head conductivity frequency response in vivo with optimized EIT-EEG. Neuroimage. 2015;

42. Janssen AM, Rampersad SM, Lucka F, Lanfer B, Lew S, Aydin U, et al. The influence of sulcus width on simulated electric fields induced by transcranial magnetic stimulation. Phys Med Biol. 2013;58: 4881–4896.

43. Heller L, van Hulsteyn DB. Brain stimulation using electromagnetic sources: theoretical aspects. Biophys J. 1992;63: 129–138.

44. Nummenmaa A, Stenroos M, Ilmoniemi RJ, Okada YC, Hamalainen MS, Raij T. Comparison of spherical and realistically shaped boundary element head models for transcranial magnetic stimulation navigation. Clin Neurophysiol. 2013;124: 1995–2007.

